# Magnetic Resonance Spectroscopy Frequency and Phase Correction Using Convolutional Neural Networks

**DOI:** 10.1101/2021.05.29.446254

**Authors:** David J. Ma, Hortense A-M. Le, Yuming Ye, Andrew F. Laine, Jeffery A. Lieberman, Douglas L. Rothman, Scott A. Small, Jia Guo

**Author notes:** Correspondence: Jia Guo.

## Abstract

We introduce DeepSPEC, a novel convolutional neural network (CNN) -based approach for frequency-and-phase correction (FPC) of MRS spectra to achieve fast and accurate FPC of single-voxel PRESS MRS and MEGA-PRESS data. In DeepSPEC, two neural networks, including one for frequency correction and one for phase correction were trained and validated using published simulated and *in vivo* PRESS and MEGA-PRESS MRS dataset with wide-range artificial frequency and phase offsets applied. DeepSPEC was subsequently tested and compared to the current deep learning solution - a “vanilla” neural network approach using multilayer perceptrons (MLP). Furthermore, random noise was added to the original simulated dataset to further investigate the model performance with noise at varied signal-to-noise (SNR) levels (i.e., 6 dB, 3 dB, and 1.5 dB). The testing showed that DeepSPEC is more robust to noise compared to the MLP-based approach due to having a smaller absolute error in both frequency and phase offset prediction. The DeepSPEC framework was capable of correcting frequency offset with 0.01±0.01 Hz and phase offset with 0.12±0.09° absolute errors on average for unseen simulated data at a high SNR (12 dB) and correcting frequency offset with 0.01±0.02 Hz and phase offset within -0.07±0.44° absolute errors on average at very low SNR (1.5 dB). Furthermore, additional frequency and phase offsets (i.e., small, moderate, large) were applied to the *in vivo* dataset, and DeepSPEC demonstrated better performance for FPC when compared to the MLP-based approach. Results also show DeepSPEC has superior performance than the model-based SR implementation (mSR) in FPC by having higher accuracy in a wider range of additional offsets. These results represent a proof of concept for the use of CNNs for preprocessing MRS data and demonstrate that DeepSPEC accurately predicts frequency and phase offsets at varying noise levels with state-of-the-art performance.

## Introduction

MRS is a widely available approach for research and for routine clinical applications, which provides noninvasive, quantitative metabolite profiles of tissue. However, many instabilities such as scanner frequency drift and subject motion can result in frequency and phase shifts that affect data analysis. Thus, frequency and phase correction (FPC) is an important technique that needs to be performed for metabolite quantification. For instance, the metabolite GABA is the primary inhibitory neurotransmitter in the human brain. A variety of studies of neurological and psychiatric disorders have shown their unique pathological characteristic in brain dysfunction [1,2]. Among a wide range of methods for measuring GABA *in vivo*, MEGA-PRESS is currently the most widely used MRS technique [3,6]. MEGA-PRESS is a J-difference editing (JDE) pulse sequence that separates GABA from overlapping metabolites such as creatine (Cr), which is present in much greater concentrations. A major limitation in JDE pulse sequences is that they depend on the subtraction of spectral edited “On Spectra” and non-edited “Off Spectra” to reveal the edited resonance in the “Diff Spectra”. As a result of the overlapping resonances being an order of magnitude larger in intensity than the GABA resonance, small changes in scanner frequency and spectral phase will lead to incomplete subtraction and distortion of the edited spectrum. The standard approach in GABA editing is to apply frequency and phase drift correction of individual frequency domain transients [4,5] by fitting the Cr signal at 3 ppm. The major limitation of the Cr fitting-based correction method is that it relies strongly on sufficient SNR of the Cr signal in the spectrum. To overcome this limitation, spectral registration (SR) approaches were proposed that can accurately align single transients in the time domain [6,7] or frequency domain [8]. In the SR approach, the frequency and phase offsets are estimated based on a nonlinear optimization numerical method to maximize the cross-correlation between each transient to a reference template. The correction accuracy depends on the overall spectral SNR. The performance of the SR method for drift correction is limited at low SNR around 1.5 dB when the spectra are dominated by noise.

Deep learning is a common strategy to address wide range of complex computational problems. The architecture of the network in terms of the number of layers, type of layers, and output function is fixed prior to inputting data. The output data is unknown, and sufficient network training is required to optimize the weights and bias parameters for assessment on validated and tested data. Moreover, deep learning is an effective image processing approach that has been enthusiastically adopted in MR imaging but thus far has had a more modest impact on MRS. A multilayer perceptron (MLP) model has been recently applied to single-transient FPC for edited MRS [9]. This work demonstrated the great potential of applying deep learning in MRS data preprocessing by pre-training models with simulated datasets with different frequency and phase offsets. Although MLP provides quick and yields well-predicted results, a more robust network could be considered, such as a convolutional neural network (CNN) approach to more accurately obtain spatial information and extract key features of the input data for FPC.

In the present study, we aim to investigate the feasibility and utility of CNNs for FPC of single voxel MEGAPRESS MRS data. We implemented automatic FPC with CNNs for the first time. Our proposed approach, DeepSPEC, was tested on a published simulated dataset and an *in vivo* dataset against the benchmark - a “vanilla” neural network approach using MLP [9]. DeepSPEC achieved state-of-the-art performance and nearly optimal correction efficiency. We then investigated the effect of additional noise of 6 dB, 3 dB, and 1.5 dB on the FPC performance to further demonstrate that DeepSPEC is a more robust solution when dealing with spectra with a low signal-to-noise ratio (SNR). Additional offsets (small, moderate, large) were also applied to the *in vivo* dataset to demonstrate the utility of DeepSPEC to accurately predict the spectral frequency and phase offsets in a more real-life scenario.

## Methods

### A. Datasets

#### 1) Simulated Datasets

For adequate network training, data selection is a vital challenge for deep learning. Since there is no ground truth of frequency and phase offsets for the *in vivo* dataset, in this work, the MEGA-PRESS training, validation, and test transients were simulated using the FID-A toolbox (version 1.2), with the same parameters as described in the previous work [9]. The training set for the DeepSPEC model was allocated 36,000 OFF+ON spectra, the validation set was allocated 4,000, and 1,000 for the test set. Furthermore, we created additional spectra with lower SNRs (at 6 dB, 3 dB, and 1.5 dB) by adding random Gaussian noise to the published simulated dataset respectively.

#### 2) In vivo Datasets

*In vivo* data was retrieved from the publicly available Big GABA repository [10]. Thirty-three MEGA-edited datasets were collected in total. 320 transients OFF+ON were used and tested on DeepSPEC, all of which were acquired using a water suppression method (VAPOR) [18] that generated positive water residual in the spectra.

### B. Network architecture

#### 1) DeepSPEC

A CNN model was evaluated to compare its accuracy in frequency and phase offset prediction (Figure 1A). The model was implemented as sequential networks (F-model, then P-model). Each network consisted of a channel with 1024 nodes as an input layer and a one-dimensional convolutional layer followed by a one-dimensional max-pooling layer. The latter layer was subsequently connected to another series of one-dimensional convolutional layer followed by a one-dimensional max-pooling layer. The convolutional layer consisted of 4 kernels with a size of 3, and the max-pooling layer has a pool size of 2 with a stride of 2. Furthermore, a fully-connected layer (FC) with 1024, 512 were used a final fully-connected linear output layer of 1 node was designed. Each hidden layer was followed by a rectified linear unit activation function to introduce non-linearity. An Adam optimizer was used to train the neural network. The output from each network was the predicted offset of frequency or phase. Adam was used as the optimizer for the DeepSPEC network with a 0.01 learning rate, and each model was trained for 300 epochs with a batch size of 32, and the mean absolute error was the loss function.

**Fig. 1.**
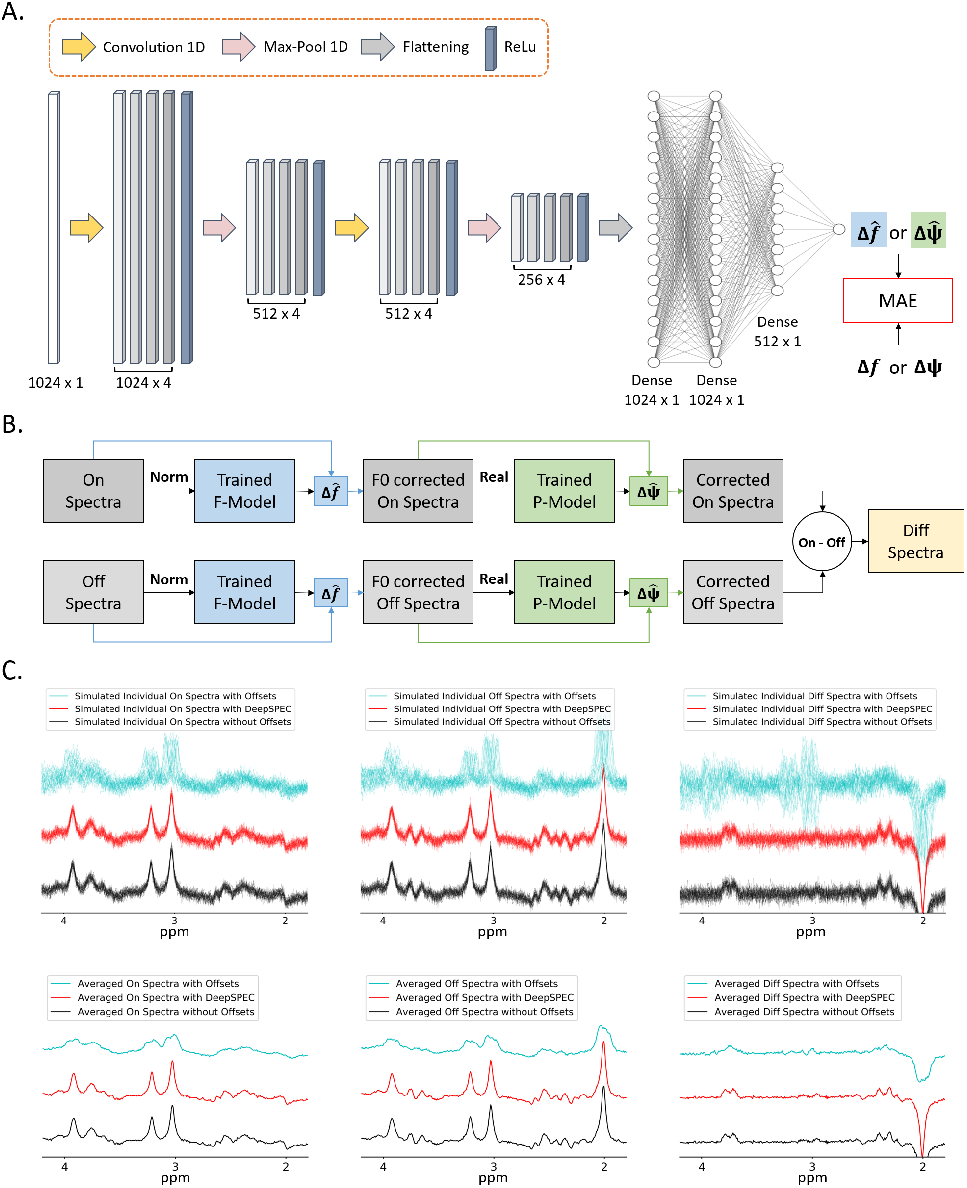
Network structure, the pipeline for assessment and sample outputs of DeepSPEC model. (A) The network architecture of the CNN model. Both the frequency and phase offset were predicted using the same architecture of 2 hidden 1D convolutional layers, 2 1D max-pooling layers, and 3 fully connected layers. The convolutional layer consisted of 4 kernels with a size of 3, and the max-pooling layer had a pool size of 2 with a stride of 2. Furthermore, two fully-connected layers (FC) with 1024 and 512 nodes respectively followed by a final fully-connected linear output layer of 1 node, were implemented. All hidden layers were each followed by a rectified linear unit (ReLU) activation function and the output fully connected layer by a linear activation function that generated the predicted offset. Simulated spectra manipulated from FID-A with artificially generated frequency or phase offsets were used as training data for the network (F-model and P-model). Each network was trained through 300 epochs with early stopping implemented when 40 consecutive epochs did not improve the lowest validation loss. (B) Flow chart of computation to determine the Diff spectra with details of the input and output from the network architecture. (C) Comparison between On, Off, and Diff spectra with offsets, with DeepSPEC, and without offsets. The top row shows multiple simulated individual spectra, while the bottom row shows the averaged spectra.

### C. Network testing

On the scale of -20 to 20 Hz and 90° to 90°, uniformly distributed artificial offsets were first added into random pairs, consisting of a frequency offset and a phase offset. Gaussian distributed noise was added to the dataset right before inputting into the network. The random pairs were then applied to the time-domain simulated transient. Figure 1B demonstrates the mechanism of DeepSPEC. The manipulation of the CNN networks for ON and OFF spectra was the same. First, we applied a Fast Fourier transform to the uncorrected data and normalized them to the maximum signal in the spectrum. The peripheral 1024 samples were then cropped off, and the central 1024 samples were selected and the absolute value was taken to feed the network. Subsequently, the predicted frequency offset (Δ*f*) was applied to the original transient to perform frequency correction. Next, we applied a Fast Fourier transform and normalized the frequency-corrected transient in the time domain. We selected and took the absolute value of the central 1024 samples into the phase-offset-prediction network. The network predicted the phase offset (Δ*ϕ*) and was used for phase correction for the frequency-corrected transient. Finally, by subtracting the corrected OFF transients from the ON transients, an average difference spectrum was obtained.

#### 1) Evaluation and comparison using in vivo dataset

The thirty-three MEGA-edited datasets were used as the test set of the DeepSPEC network. For a first comparison to the performance of DeepSPEC CNN, SR [14] performed FPC in the time domain. ON and OFF transients were registered to a single template, and the first *n* points of the signal were used, where n was the last point at which the SNR was higher than 3. The noise was computed from the bottom quarter of the signal, and n was set to a value larger than 100. The real and imaginary parts of the first n points were concatenated as a real vector and registered to the median transient of the dataset using MatLab function nlinfit (version 2019a, MathWorks, Natick, MA). The starting parameters for the subsequent transient were the same as the fitted parameters from 1 transient. The initial starting values for the offsets were 0 Hz and 0 degrees. In order to correct for the residual frequency and phase offsets, the transients were averaged, and global FPC was performed using Cr/Cho modeling (nlinfit) of this averaged spectrum after registration. Beyond SR, a model-based SR (mSR) was also implemented as a comparison of DeepSPEC. Unlike SR, mSR uses a noise-free model as the template instead of the median transient of the dataset. Noise-free ON and OFF FID models were created in Osprey (version 1.0.0),[12] an open-source MatLab toolbox, following peer-reviewed preprocessing recommendations.[11] As another comparison for DeepSPEC CNN, a “vanilla” neural network using MLP containing 3 FC layers (1024, 512, 1 node(s)) was tested.[9] In this network, each hidden FC layer was followed by a ReLU activation function, and a linear activation function followed the output layer.

To test the network in a more extreme environment, in addition to the random offsets, additional artificial offsets were added to the *in vivo* data. There were 3 different kinds of additionally added offsets: 1. 0 ≤|Δ*f* | ≤ 5*Hz* and 0 ≤ | Δ*ϕ*| ≤ 20°; 2. 5≤|Δ*f*|≤10*Hz* and 20° ≤|Δ*ϕ*|≤45°; 3. 10 ≤ |Δ*f*|≤20*Hz* and 45° ≤ |Δ*ϕ* | ≤90°. All additional offsets were sampled from a uniform distribution and added as random pairs of frequency and phase to each transient.

### D. Performance measurement

In the simulated dataset, the artificial offsets were set as the ground truth, and the mean square error between the ground truth and predicted value was used as the criteria to measure the network’s performance. Moreover, we calculated and plotted the difference value between the true spectra and the corrected spectra using SR, MLP, and DeepSPEC. A Q score was used to determine the performance strengths of each different methods, and it is defined as: *Q* = 1 − *σ*_12_*/*(*σ*_12_ +*σ*_22_), where *σ*_2_ is the variance of the choline subtracted artifact in the average difference spectrum. If the Q score is greater than 0.5, it indicates that the first method performs better than the second method and vice versa.

## Results

### A. Spectra Analysis between the MLP-based approach and DeepSPEC for varying SNRs

Figures 2A, 2C, 2E, and 2G illustrate the results of the MLP-based approach on the 1000 transients (500 ON, 500 OFF) of the simulated test set as well as of the test set with lower SNR at 6 dB, 3 dB, and 1.5 dB. Figures 2B, 2D, 2F, and 2H show the results of the DeepSPEC on the same simulated test set as well as of the test set with lower SNR at 6 dB, 3 dB, and 1.5 dB. In each subfigure of Figure 2, the frequency offset errors are plotted against their corresponding correct values, the phase offset errors are plotted against their corresponding correct values, the model-corrected difference spectrum and the difference spectrum corrected by the true offsets are plotted together, and the residues between the difference spectra are shown. For the original test set, the mean frequency offset error was 0.02 ± 0.02 Hz for the MLP-based approach and 0.01 ± 0.01 Hz for the DeepSPEC, and the mean phase offset error was 0.19 ± 0.17° for the MLP-based approach and 0.12 ± 0.09° for the DeepSPEC. With a lower SNR at 6 dB, the mean frequency offset error was 0.00 ± 0.04 Hz for the MLP-based approach and 0.00 ± 0.02 Hz for the DeepSPEC, and the mean phase offset error was 0.02 ± 0.36° for the MLP-based approach and -0.08 ± 0.29° for the DeepSPEC. With a lower SNR at 3 dB, the mean frequency offset error was 0.00 ± 0.05 Hz for the MLP-based approach and -0.01 ± 0.02 Hz for the DeepSPEC, and the mean phase offset error was 0.01 ± 0.41° for the MLP-based approach and 0.01 ± 0.34° for the DeepSPEC. With a lower SNR at 1.5 dB, the mean frequency offset error was 0.00 ± 0.05 Hz for the MLP-based approach and 0.01 ± 0.02 Hz for the DeepSPEC, and the mean phase offset error was 0.02 ± 0.61° for the MLP-based approach and -0.07 ± 0.44° for the DeepSPEC. Figure 2 shows that the DeepSPEC had smaller errors within the frequency and phase ranges for all SNR levels tested.

**Fig. 2.**
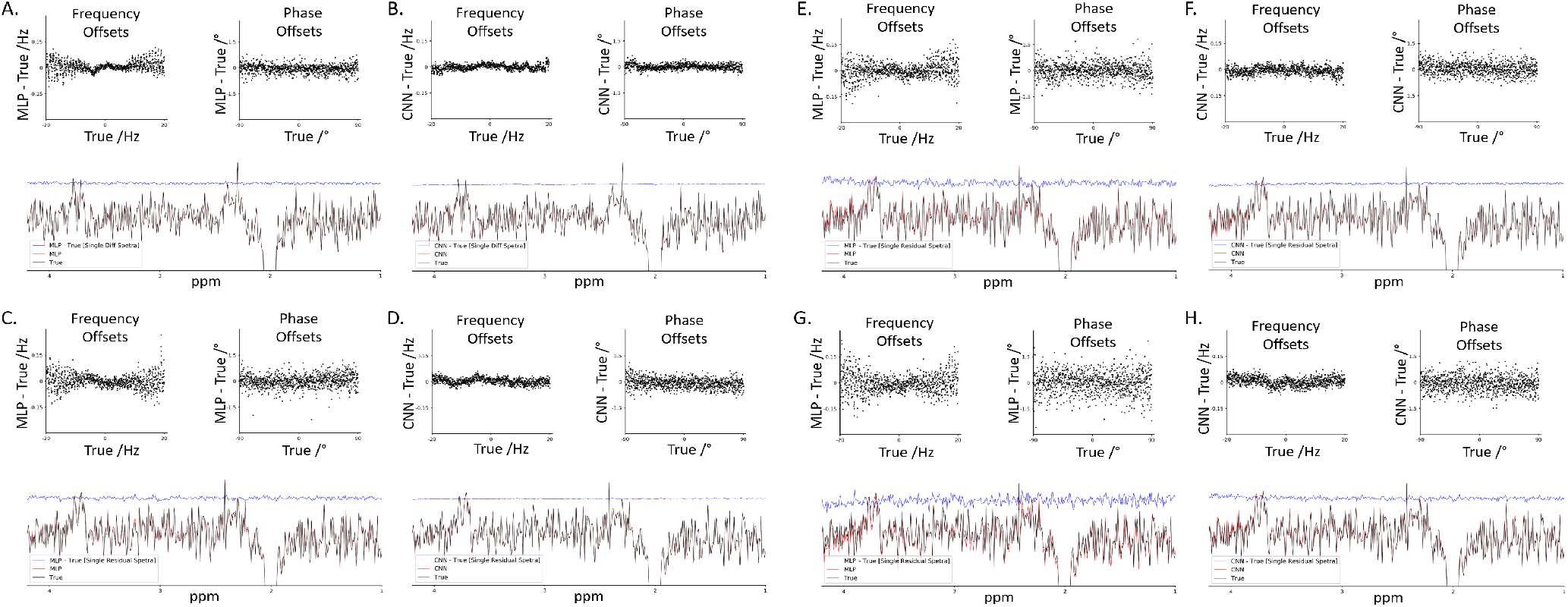
Visualizing the performance of the MLP-based approach and our proposed DeepSPEC for frequency and phase correction using the published simulated dataset for varying SNRs. The scatter plots show the correction errors between the ground truths and model predictions at different frequency and phase offsets. The spectra below the scatter plots demonstrate the deep learning model predictions (MLP or CNN), the true MEGA-PRESS difference spectrum, and the subtraction of the MLP/CNN and true difference spectrum (Single Diff Spectra). (A) Output of MLP-based approach on the original test set (SNR at 12 dB). (B) Output of DeepSPEC on the original test set. (C) Output of MLP-based approach on the test set with SNR at 6 dB. (D) Output of DeepSPEC results on the test set with SNR at 6 dB. (E) Output of MLP-based approach on the test set with SNR at 3dB. (F) Output of DeepSPEC on the test set with SNR at 3 dB. (G) Output of MLP-based approach on the test set with SNR at 1.5 dB. (H) Output of DeepSPEC on the test set with SNR at 1.5 dB.

### B. DeepSPEC and MLP-based approach Error comparison

Figure 3 illustrates the comparison of the results of the MLP-based approach and the DeepSPEC for frequency and phase correction of both the Off spectra and the On spectra of the simulated test set for varying SNRs (high, medium, low). DeepSPEC showed significantly lower frequency estimation errors than the MLP-based model for the Off spectra at varying SNRs (Figure 3C) and for the On spectra at varying SNRs (Figure 3A). For example, with the test set at a lower SNR of 1.5 dB, the mean frequency offset error for Off spectra was 0.042 ± 0.036 Hz for the MLP-based approach and 0.019 ± 0.015 Hz for the DeepSPEC, and for On spectra, it was 0.041 ± 0.036 Hz for the MLP-based approach and 0.021 ± 0.016 Hz for the DeepSPEC (Figure 3H). Similarly, it showed significantly lower phase estimation errors than the MLP-based model for the OFF spectra at varying SNRs (Figure 3D) and for the On spectra at varying SNRs (Figure 3B). With the test set at a lower SNR of 1.5 dB, the mean phase offset error for Off spectra was 0.429 ± 0.351° for the MLP-based approach and 0.372 ± 0.289° for the DeepSPEC, and for On spectra it was 0.518 ± 0.436° for the MLP-based approach and 0.333 ± 0.247° for the DeepSPEC (Figure 3H). Consequently, the residual spectra errors of the DeepSPEC were significantly lower than those of the MLP-based model for the Off spectra at varying SNRs (Figure 3F), for the On spectra at varying SNRs (Figure 3E), and for the difference spectra between the MLP-based model and the DeepSPEC at varying SNRs (Figure 3G), indicating the overall higher performance of the DeepSPEC with respect to the MLP-based approach as well as its robustness to noise.

**Fig. 3.**
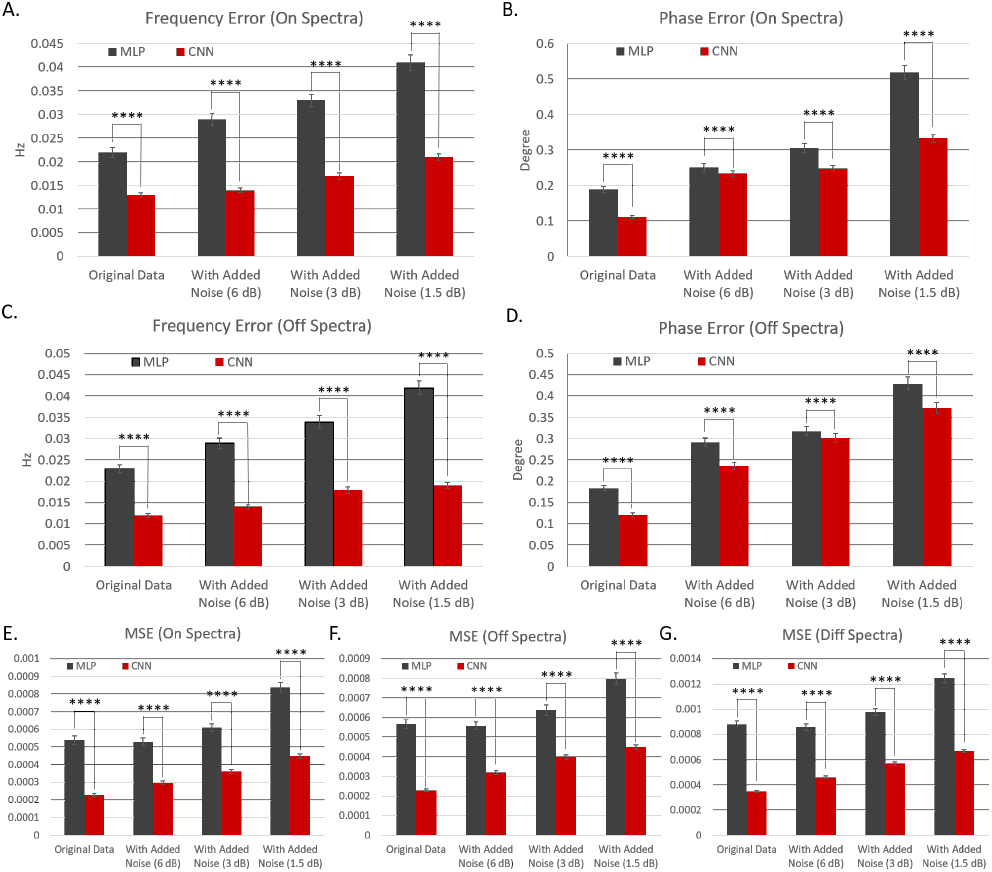
Comparison between the MLP-based approach and our proposed DeepSPEC for frequency and phase correction of both the OFF spectra and On spectra at varying SNRs. (A) Bar graph showing the frequency estimation error (in Hz) of the MLP-based method and the DeepSPEC at varying SNRs of the On spectra. (B) Bar graph showing the phase estimation error (in degrees) of the MLP-based method and the DeepSPEC method at varying SNRs of the On spectra. (C) Bar graph showing the frequency estimation error (in Hz) of the MLP-based method and the DeepSPEC at varying SNRs of the Off spectra. (D) Bar graph showing the phase estimation error (in degrees) of the MLP-based method and the DeepSPEC method at varying SNRs of the Off spectra. (E) Bar graph showing the residual spectra mean squared error of the MLP-based method and the DeepSPEC of the On spectra. (F) Bar graph showing the residual spectra mean squared error of the MLP-based method and the DeepSPEC of the Off spectra. (G) Bar graph showing the residual spectra error of the MLP-based method and the DeepSPEC of the difference spectra. The two-tailed P value was used and is less than 0.0001**** for all the comparisons between the MLP-based and DeepSPEC-based approaches.

As observed in Figures 2 and 3, the original test set’s results were fairly better than the results of the test set with added noise of 6 dB, indicating that the noise level in the original test set must have been higher than 6 dB. Further computations confirmed that the test set’s noise level was 12 dB.

**Table 1.**
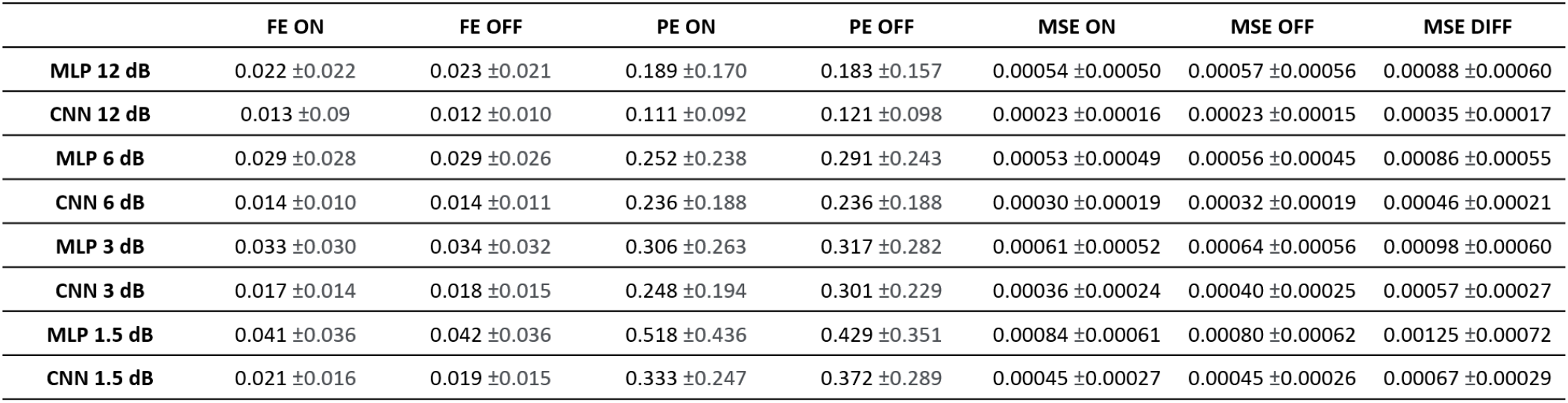
Table of the errors of the MLP-based approach and our proposed DeepSPEC for frequency and phase correction of the On spectra and Off spectra, of the resulting On and Off spectra, and of the corresponding difference spectra at varying SNRs.

### C. in vivo Datasets

Figure 4A illustrates the spectra resulting from the 33 *in vivo* datasets without (column 1) or with additional artificial offsets (columns 2-4) for no correction, MLP-based correction, DeepSPEC-based correction, and SR-based correction. When small offsets C1 were added, all three models performed similarly. The mean performance score of the MLP-based approach and DeepSPEC was 0.51 ± 0.09 (Figure 4D, column 2) while it was 0.49 ± 0.08 for DeepSPEC and SR (Figure 4C, column 2), and 0.49 ± 0.10 for the MLP-based approach and SR (Figure 4B, column 2). the MLP-based approach performed better than SR for 42.42 % of the 33 *in vivo* datasets, DeepSPEC performed better than SR for 45.45% of the 33 *in vivo* datasets, and DeepSPEC performed better than the MLP-based approach for 66.67 % of the 33 *in vivo* datasets. As for medium offsets C2, the performance of DeepSPEC and the MLP-based approach was comparable, but both models performed better than SR. The mean performance score of the MLP-based approach and DeepSPEC was 0.54 ± 0.09 (Figure 4D, column 3) while it was 0.78± 0.14 for DeepSPEC and SR (Figure 4C, column 3), and 0.79 ± 0.13 for the MLP-based approach and SR (Figure 4B, column 3), the MLP-based approach performed better than SR for 96.97 % of the 33 *in vivo* datasets, DeepSPEC performed better than SR for 96.97 % of the 33 *in vivo* datasets, and DeepSPEC performed better than the MLP-based approach for 60.61 % of the 33 *in vivo* datasets. When large offsets C3 were added, the performance of DeepSPEC was better than the MLP-based approach and SR’s. The mean performance score of the MLP-based approach and DeepSPEC was 0.57 ± 0.14 (Figure 4D, column 4) while it was 0.77 ± 0.12 for DeepSPEC and SR (Figure 4C, column 4), and 0.73 ± 0.16 for the MLP-based approach and SR (Figure 4B, column 4). The MLP-based approach performed better than SR for 90.91 % of the 33 *in vivo* datasets, DeepSPEC performed better than SR for 96.97 % of the 33 *in vivo* datasets, and DeepSPEC performed better than the MLP-based approach for 75.76 % of the 33 *in vivo* datasets. For small and medium offsets, DeepSPEC-corrected spectra and MLP-corrected spectra (Figure 4A, columns 2-3) are similar to the original spectra (Figure 4A, column 1). However, for large offsets, the MLP-corrected spectra (Figure 4A, column 4) slightly diverge from the original spectra, while the DeepSPEC-corrected spectra still are not noticeably different from the original spectra. mSR exhibited the same performance pattern as DeepSPEC with respect to the MLP-based approach, with a similar mean performance score of 0.51 ± 0.13 for small offsets and of 0.52 ± 0.13 for medium offsets, and a better mean performance score of 0.57 ± 0.11 for large offsets. (All numerical results are shown in Figure 4E).

**Fig. 4.**
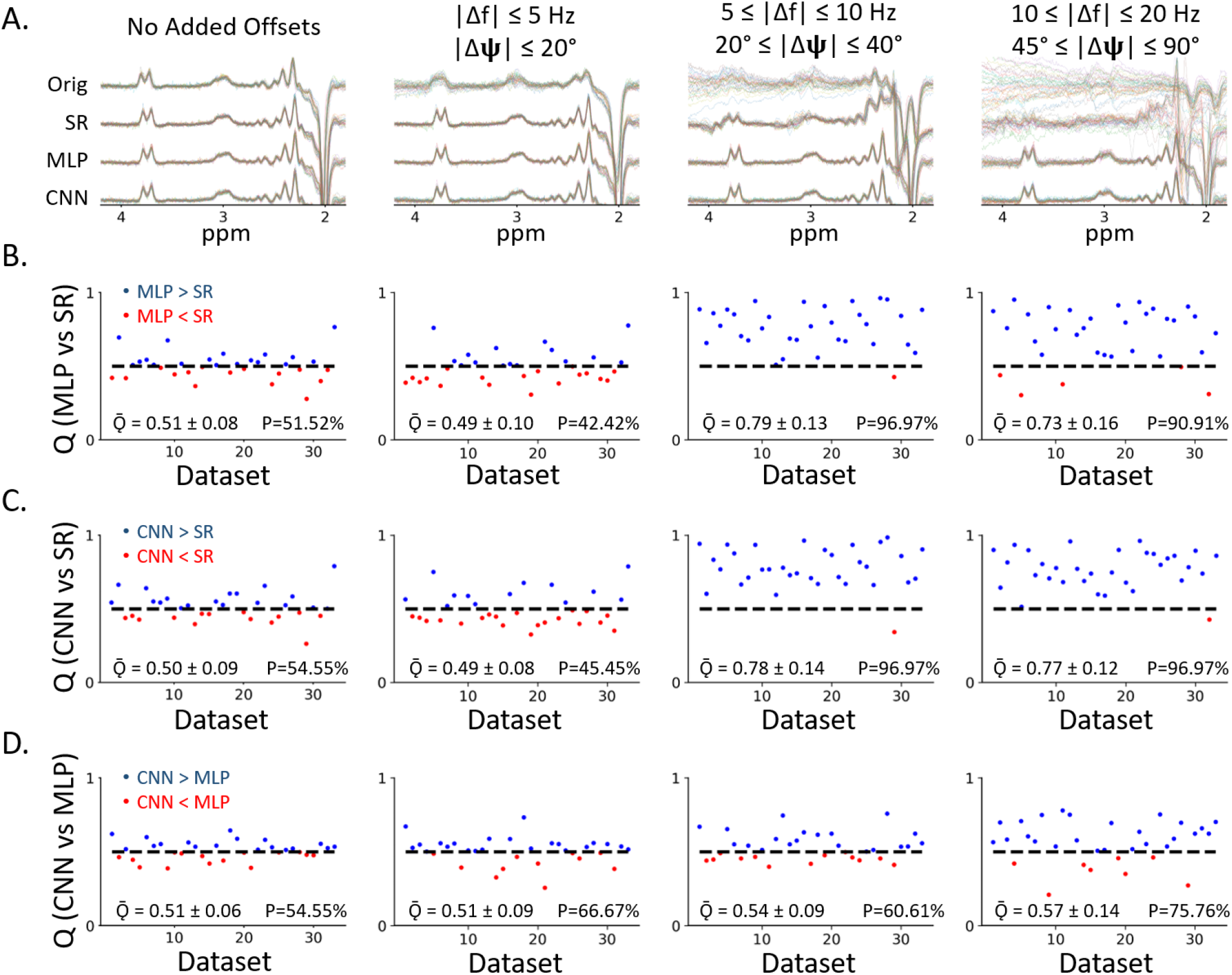
Difference spectra and performance scores comparing the MLP-based approach to SR, DeepSPEC to SR and DeepSPEC to the MLP-based approach for the 33 *in vivo* datasets. The difference spectra were generated without any FPC (NO), using the network shown in Figure 1 (MLP-based approach and CNN) and using SR (spectral registration). (A) Results of applying corrections to the *in vivo* data without further manipulation, and with additional frequency and phase offsets, applied to the same 33 datasets: small offsets (0-5 Hz; 0-20°), medium offsets (5-10 Hz; 20-45°), and large offsets (10-20 Hz; 45-90°). (B) Comparative performance scores P and Q for the MLP-based approach and SR for each dataset. A score above 0.5 indicated that the MLP-based approach performed better than SR, whereas a score below 0.5 (50%) indicated that SR performed better than the MLP-based approach in terms of alignment. (C) Comparative performance scores P and Q for CNN and SR for each dataset. (D) Comparative performance scores P and Q for CNN and the MLP-based approach for each dataset.

## Discussion

Frequency and phase correction is a crucial step when quantifying metabolites to analyze edited MRS data. The resulting Diff spectra must be as robust as possible to achieve the best result for quantification. Thus, when constructing the network architecture, meticulous attention was given to generate an optimal model. Many methodological options were considered, and we decided that having separate networks to perform frequency and phase correction using a convolutional neural network would produce the best results. A CNN model, DeepSPEC, was constructed with 2 layers of convolutional layers followed by 2 layers of max-pooling layers with an activation function of ReLU on top of the MLP-based model. Furthermore, the networks were designed to output the FPC parameters instead of the corrected spectra. Inputs were the magnitude spectrum for the frequency network and the real spectrum for the phase network. The networks were trained using simulated datasets concatenated with Off and On transients and validated using simulated datasets with artificially added offsets as well. The risk of noise was also considered where the central 1024 points were selected for the input. The robustness of the models to noise was also quantified where the noise of 6 dB, 3 dB, and 1.5 dB were introduced to the simulated dataset and compared to the MLP-based model. *In vivo* data were also used to assess the performance of the models. With given data and varied additional offsets, the performance score P and its average score Q were used. Comparison of DeepSPEC was performed with MLP, SR, and mSR for the *in vivo* datasets.

From Figure 2 and 3, we observed that the DeepSPEC model performed better compared to MLP-based models when using simulated data. The scatter plot shows the CNN model has smaller correction errors (i.e., smaller mean absolute error and standard deviation) for both frequency and phase offset estimation compared to the MLP-based approach. Moreover, it can also be seen that for each offset, the subtraction of the prediction to the ground truth data points congregate near the *y=0* line indicating the small deviation it had compared to the true value. In addition, the Diff spectra below also demonstrated that our model has smaller residual differences when the corrected spectra are subtracted from the truth spectra. The influence of noise was also found to have smaller effects in predicting the Diff spectra as smaller residues were observed when the spectra were subtracted. Especially at lower SNRs (1.5 and 3 dB), the difference was obvious, indicating that the MLP-based approach cannot cope with noisy data very well. From Figure 2, a prediction of the innate SNR for the original dataset can be made. By comparing Figure 2A with other figures, it can be seen that 2B and 2D resemble each other really well. Figure 2B shows a flat residual curve indicating the SNR could be greater than 6 dB and close to 12 dB. Computation was done to confirm that the results where using the mean of the central data points of one of the truth spectra as the signal and the standard deviation of the remaining points as the noise; the SNR was computed to be approximately 12 dB. Furthermore, Figure 3 shows the mean absolute error to be smaller in all cases for DeepSPEC with improved accuracy in larger offsets when compared to the MLP-based approach. *This result can be seen for frequency, phase, and residual error where the two-tailed p-value shows the significance of this result*. Especially in frequency (On and Off) and residual error (On, Off, Diff), the differences are tremendous.

From Figure 4, the same conclusion can be made when using *in vivo* data. There is better performance in the DeepSPEC model as performance score P is higher and the output spectra are clearer compared to the MLP-based approach. This is well shown in the performance when we added a large offset to the dataset, indicating our model can handle more varied offsets with higher accuracy. The benchmark was the comparison to SR, where both MLP and CNN performed better with no offset, C1, and C3 which show superior results for DeepSPEC due to its larger Q and P-value. When comparing DeepSPEC to MLP directly (Figure 4D), it can be shown that our model performed better than MLP in all cases for both performance score P and its Q value. Additionally, comparing the performance to mSR, it can be stated that our model performs better due to the larger score for C1 and C2, indicating the lack of performance for this small and moderate offset for mSR. Smaller standard deviation is also observed throughout our DeepSPEC model performance which corresponds to the higher robustness of our model.

DeepSPEC was found to demonstrate accurate quantification with training and validation for frequency and phase offset estimation in separate models. However, this takes more computational time to train the models and makes the method inflexible to account for additional parameters such as first-order phase, amplitude, and bandwidth variance in different transients. Moreover, only the MEGA-PRESS sequence was considered in this study. The DeepSPEC performance on other JDE sequences, such as MEGA-sLASER [17] was not discussed. The magnetic field strength to produce the simulated dataset and to acquire the *in vivo* dataset was 3 T. However, the DeepSPEC performance over higher magnetic fields is yet unknown. Additionally, our study was only based on human MRS data. Nevertheless, in pre-clinical studies, animal models continue to play a significant role in the scientific investigation of neuropsychiatric disorders. Future studies on animal MRS data may further validate the generalization of the proposed method across species. Data from different vendors other than Philips such as Siemens, GE, and Bruker are also an important variable that should be considered in the future. Finally, different experimental conditions such as *ex vivo, in situ*, and *in vitro* should also be assessed to further demonstrate our model’s general utilities in MRS data preprocessing.

For future work, combining frequency and phase offset predictions into a single model can further improve the computational efficiency. Although DeepSPEC was implemented and optimized specifically for MEGA-PRESS MRS data at 3T in this study, DeepSPEC should be able to process any MRS spectra in general. Application-specific MRS training data across different sequences, field strengths, and varying experimental conditions can be simulated using publicly available toolboxes such as SpinWizard [15], FID-A, and MARSS [16], and the tailored DeepSPEC networks can be easily retrained for any MRS experiment accordingly.

**Table 2.** Table of the P score percentages and the average Q scores comparing the MLP-based approach to SR, CNN to SR and CNN to the MLP-based approach with no additional offsets and three magnitudes of additional offsets (C1. 0 ≤ |Δ*f*| ≤ 5*Hz* and 0 ≤ |Δ*ϕ*| ≤ 20°; C2. 5 ≤ |Δ*f*| ≤ 10*Hz* and 20° ≤ |Δ*ϕ*| ≤ 45°; C3. 10 ≤ |Δ*f*| ≤ 20*Hz* and 45° ≤ |Δ*ϕ*| ≤ 90°). P score percentages and the average Q score of mSR to MLP are also depicted.

|  | No Added Offsets | C1 | C2 | C3 |
| --- | --- | --- | --- | --- |
| MLP vs. SR (%) | 51.52% | 42.42% | 96.97% | 90.91% |
| MLP vs. SR (Q) | 0.51 ± 0.08 | 0.49 ± 0.10 | 0.79 ± 0.13 | 0.73 ± 0.16 |
| CNN vs. SR (%) | 54.55% | 45.45% | 96.97% | 96.97% |
| CNN vs. SR (Q) | 0.50 ± 0.09 | 0.49 ± 0.08 | 0.78 ± 0.14 | 0.77 ± 0.12 |
| CNN vs. MLP (%) | 54.55% | 66.67% | 60.61% | 75.76% |
| CNN vs. MLP (Q) | 0.51 ± 0.06 | 0.51 ± 0.09 | 0.54 ± 0.09 | 0.57 ± 0.14 |
| mSR vs. MLP (%) | 54.55% | 48.48% | 57.58% | 75.76% |
| mSR vs. MLP (Q) | 0.51 ± 0.12 | 0.51 ± 0.13 | 0.52 ± 0.13 | 0.57 ± 0.11 |

## Conclusions

This work provides the first proof of concept of the feasibility of a CNN framework to preprocess MRS spectra. Our DeepSPEC model shows better performance and delivers results more robust to noise as compared to the current state-of-the-art model in both simulation and *in vivo* tests.

## Acknowledgements

This study was supported by the Technical Development Grant Program at the Columbia MR Research Center. This study was performed at the Zuckerman Mind Brain Behavior Institute MRI Platform, a shared resource.

